# Chromatin profiling of the repetitive and non-repetitive genome of the human fungal pathogen Candida albicans

**DOI:** 10.1101/475996

**Authors:** Robert Jordan Price, Esther Weindling, Judith Berman, Alessia Buscaino

## Abstract

**Background:** Eukaryotic genomes are packaged into chromatin structures with pivotal roles in regulating all DNA-associated processes. Post-translational modifications of histone proteins modulate chromatin structure leading to rapid, reversible regulation of gene expression and genome stability which are key steps in environmental adaptation. *Candida albicans* is the leading fungal pathogen in humans, and can rapidly adapt and thrive in diverse host niches. The contribution of chromatin to *C. albican*s biology is largely unexplored.

**Results:** Here, we harnessed genome-wide sequencing approaches to generate the first comprehensive chromatin profiling of histone modifications (H3K4me^3^, H3K9Ac, H4K16Ac and γ-H2A) across the *C. albicans* genome and relate it to gene expression. We demonstrate that gene-rich non-repetitive regions are packaged in canonical euchromatin associated with histone modifications that mirror their transcriptional activity. In contrast, repetitive regions are assembled into distinct chromatin states: subtelomeric regions and the rDNA locus are assembled into canonical heterochromatin, while Major Repeat Sequences and transposons are packaged in chromatin bearing features of euchromatin and heterochromatin. Genome-wide mapping of γH2A, a marker of genome instability, allowed the identification of potential recombination-prone genomic sites. Finally, we present the first quantitative chromatin profiling in *C. albicans* to delineate the role of the chromatin modifiers Sir2 and Set1 in controlling chromatin structure and gene expression.

**Conclusions:** This study presents the first genome-wide chromatin profiling of histone modifications associated with the *C. albicans* genome. These epigenomic maps provide an invaluable resource to understand the contribution of chromatin to *C. albicans* biology.

## BACKGROUND

Packaging of genomes into chromatin is the key determinant of nuclear organization [1,2]. The basic unit of chromatin is the nucleosome, consisting of a histone octamer of two molecules each of histone H2A, H2B, H3, and H4, around which 147 bp of DNA are wrapped in almost two turns [3]. Histone proteins are subjected to a wide variety of post-translational modifications, known as histone marks, that decorate distinct chromatin regions [3]. Modification of chromatin structure controls a plethora of nuclear processes including gene expression, DNA repair and DNA replication [3,4]. Consequently, genome-wide maps of histone modifications have been instrumental in identifying functionally different regions of eukaryotic genomes [5,6]. Gene-rich, non-repetitive DNA is associated with active histone marks, forming euchromatin, a chromatin state permissive to transcription and recombination [7]. At euchromatic regions, promoters of active genes are enriched in histone H3 trimethylated on lysine 4 (H3K4me^3^) and acetylated on lysine 9 (H3K9Ac), while gene bodies are enriched in a different set of histone modifications, such as acetylation of lysine 16 on histone H4 (H4K16Ac) [8–10]. In contrast, genomic regions enriched in repetitive DNA and low in gene density are assembled into heterochromatin [7]. These repetitive sequences (including tandem repeats, transposable elements and gene families) are a threat to genome stability. At repetitive elements, heterochromatin assembly promotes genome stability by repressing deleterious recombination events [7,11,12]. Heterochromatin is devoid of active histone marks (i.e. H3K4me^3^, H3K9Ac and H4K16Ac) and is enriched in repressive histone marks such as methylation of lysine 9 on histone H3 (H3K9me) and methylation of lysine 27 on histone H3 (H3K27me) [1].

While euchromatin structure is largely conserved across organisms, histone marks associated with heterochromatic regions vary between organisms. For example, in the model system *Saccharomyces cerevisiae*, heterochromatin is devoid of H3K9me and H3K27me marks but nucleosomes are hypomethylated on H3K4 and hypoacetylated on H3K9 and H4K16 [1,13]. Phosphorylation of serine 129 on histone H2A (known as γ-H2A) is enriched at heterochromatin regions in *S. cerevisiae*, *Schizosaccharomyces pombe* and *Neurospora* crassa, independently of the cell cycle stage [14–17]. Given that γ-H2A is a hallmark of DNA double strand breaks, these findings suggest that heterochromatic regions are flagged for DNA damage. In contrast, in human cells, phosphorylation of H2AX, a modification functionally analogous to γ-H2A, does not decorate heterochromatic regions [18,19]. Chromatin modifications also play major roles in controlling genome stability by dictating pathways of DNA repair. Indeed, choices of DNA repair pathways (i.e. Non Homologous End Joining or Homologous Recombination) depend on the chromatin state of the genomic region undergoing repair and extensive chromatin changes, including γ-H2A, are linked to repair of DNA breaks [20]. Consequently, in unchallenged cells, γ-H2A mapping is used to identify unstable genomic regions (named γ-sites) that are prone to intrinsic DNA damage and recombination [14]. Chromatin modifications are reversible and specific histone modifiers maintain or erase the histone modification state associated with different chromatin regions. Among these, histone acetyltransferases (HATs) and histone deacetylases (HDACs) respectively maintain and erase histone acetylation, while histone methyltransferases (HMTs) and demethylases (HDMs) are responsible for the methylation state of histones [2,3]. Chromatin regulation rapidly and reversibly alters gene expression and genome stability and can, therefore, have a major impact on environmental adaptation of microbial organisms that need to rapidly adapt to sudden environmental changes [21,22].

One such organism is the human fungal pathogen *Candida albicans*. *C. albicans* is a commensal organism that colonises the mouth, the skin, and the uro-intestinal and reproductive tracts of most individuals without causing any harm. However, *C. albicans* is also the most common causative agent of invasive fungal infections and systemic infections are associated with high mortality rates (up to 50%) [23]. *C. albicans* is such a successful pathogen because it rapidly adapts and thrives in diverse host niches. The ability to switch among multiple specialised cell types, as well as its remarkable genome plasticity, is at the basis of *C. albicans* adaptation [24].

*C. albicans* is a diploid organism with a genome organised in 2 × 8 chromosomes containing 6408 protein-coding genes, in addition to a large number of non-coding RNAs [25–28]. The genome of *C. albicans* contains several classes of repetitive elements: telomeres/subtelomeres, the rDNA locus, Major Repeat Sequences (MRS) and transposable elements [29]. Telomeres are composed of tandemly repeating 23□bp units, while subtelomeres are enriched in long terminal repeats (LTR), retrotransposons and gene families [29,30].

The rDNA locus consists of a tandem array of a ~12□kb unit repeated 50 to 200 times. Each unit contains the two highly conserved 35□S and 5□S rRNA genes that are separated by two Non-Transcribed Spacer regions (NTS1 and NTS2), whose sequences are not conserved with other eukaryotes [29,31].

MRS loci are long tracts (10–100□kb) of nested DNA repeats found on 7 of the 8 *C. albicans* chromosomes [29,32]. These repetitive domains, found in *C. albicans* and in the closely related species *C. dubliensis* and *C. tropicalis*, are formed by large tandem arrays of 2.1□kb RPS unit flanked by non-repetitive HOK and RBP-2 elements. Each RBP-2 element contains a protein-coding gene, *FGR6,* important for morphological switches [32,33].

Several classes of retrotransposons are present in the *C. albicans* genome including 16 classes of LTR retro-transposons (Tca1-16) and Zorro non-LTR retrotransposons that are present in 5-10 copies per cell, dispersed along the chromosomes. Among those, Tca2, Tca4, Tca5, Zorro-2 and Zorro-3 are capable of transposition [34–36]. The *C. albicans* genome is remarkably plastic, and natural isolates exhibit a broad spectrum of genomic variations including Loss of Heterozygosity (LOH) events, chromosome rearrangements and aneuploidy [37]. Evolution experiments and analyses of clinical isolates have demonstrated that repetitive elements are hypermutable sites of the *C. albicans* genome and are prone to high rates of recombination [37,38].

Several studies have demonstrated that regulation of chromatin structure plays critical roles in regulating *C. albicans* gene expression and genome instability [39– 42]. However, comprehensive profiling of histone modifications across the whole *C. albicans* genome is still lacking. Generation of these epigenomic maps will be essential to truly understand the impact of chromatin regulation to *C. albicans* adaptation and development of virulence traits.

In this study, we used chromatin immunoprecipitation with massively parallel sequencing (ChIP-seq) technology to establish the first comprehensive genome-wide map of *C. albicans* histone modifications (H3K4me^3^, H3K9Ac, H4K16Ac and γH2A), marking euchromatic, heterochromatic regions and potential recombination-prone unstable sites. Genome-wide mapping of RNA Polymerase II (RNAPII) and transcriptome expression profiling allowed us to unveil the link between histone modification states and transcriptional activity. We demonstrate that specific chromatin states are associated with the repetitive and non-repetitive *C. albicans* genome. While gene-rich regions are associated with active chromatin marks mirroring their transcriptional state, different types of repetitive elements are assembled into distinct chromatin types. Finally, we present the first *C. albicans* quantitative ChIP-seq (q-ChIP-seq) methodology that has permitted us to elucidate the roles of the HDAC Sir2 and the HMT Set1 in shaping the chromatin state of *C. albicans* genome and regulating gene expression.

## RESULTS

### Genome-wide histone modification profiling in C. albicans

The *C. albicans* genome contains two homologous pairs of divergently transcribed histone H2A and H2B genes, and histone H3 and H4 genes in addition to a single histone H3 gene (Fig S1 A). Sequence alignment demonstrated that the frequently modified amino acid residues H3K4, H3K9, H4K16 and H2AS129 are conserved in *C. albicans* (Fig S1 B).

To explore the chromatin signature of *C. albicans* repetitive and non-repetitive regions, we globally mapped the genomic locations of H3K4me^3^, H3K9Ac, H4K16Ac and γH2A by performing Chromatin ImmunoPrecipitation followed by high-throughput sequencing (ChIP-seq). Since nucleosomes are not equally distributed across genomes, we accounted for nucleosome occupancy by performing genome-wide profiling of unmodified histone H3 and histone H4. Finally, to correlate specific histone modification profiling with transcriptional activity, we mapped RNA polymerase II (RNAPII) occupancy genome-wide. In parallel, we performed transcriptome analysis by strand-specific RNA sequencing (RNA-seq) to profile gene expression levels.

For all samples, ChIP-seq was performed from *C. albicans* wild-type (WT) cells grown in standard laboratory growth conditions (YPAD 30 °C) using antibodies specific for modified or unmodified histones. Input (I) and Immunoprecipitated samples (Ip) were sequenced using the Illumina HiSeq2000 platform (single-end 50 bp reads; average coverage: 28x; Table S2, Dataset S1) and aligned to a custom haploid version of Assembly 22 of the *C. albicans* genome [25]. Unmodified histone H3 occupancy showed a strong positive correlation with histone H4 occupancy (Pearson correlation coefficient r = 0.97), with the exception of centromeric regions where the histone H3 variant Cse4^CENP-A^ replaces histone H3 (Fig 1A, 1C and S2). Furthermore, RNAPII occupancy showed a positive correlation with gene expression levels (Pearson correlation coefficient r = 0.72) (Fig 1B).

**Figure 1.**
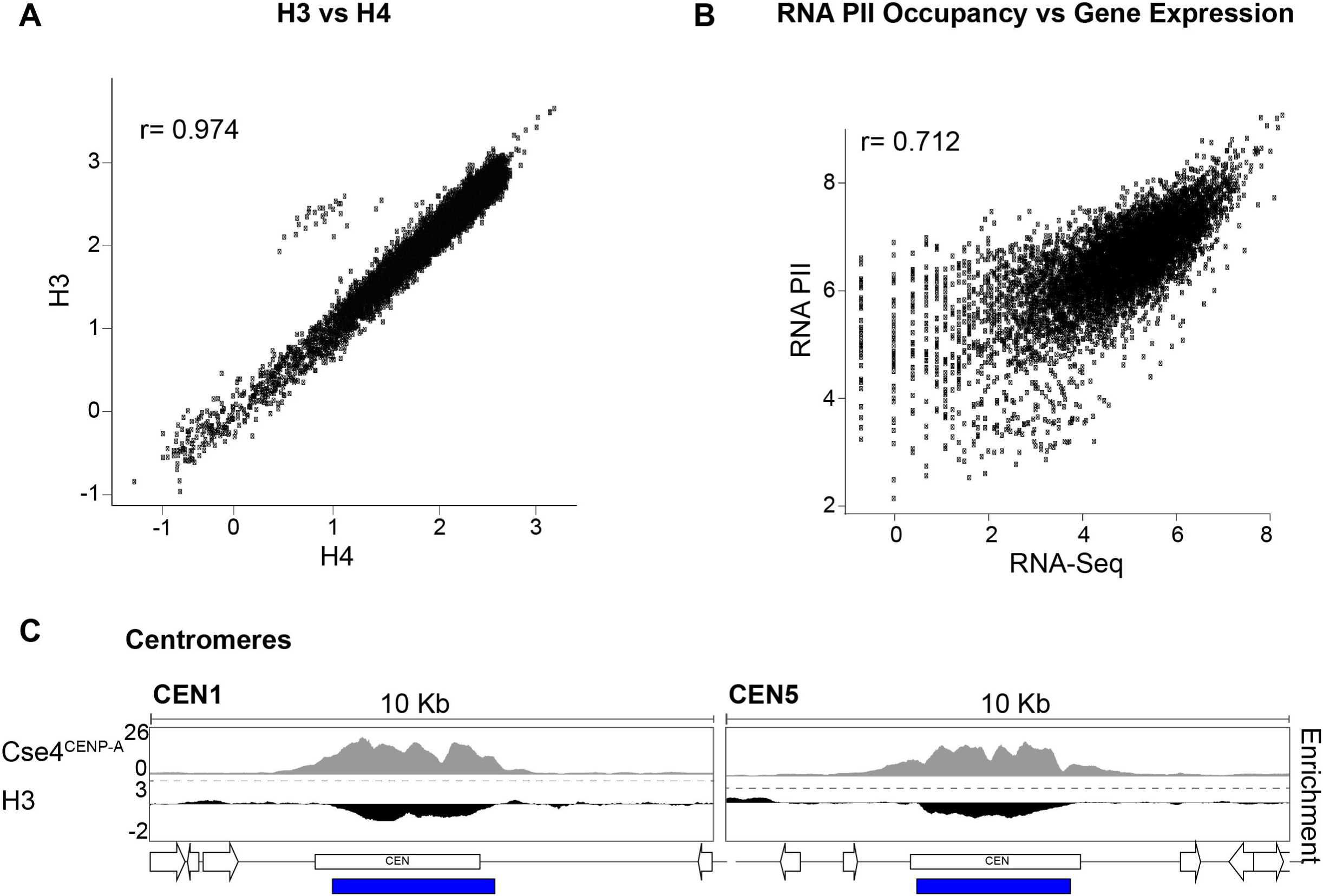
Histones and RNA Polymerase II occupancy **(A)** Correlation between H3 and H4 occupancy (log10 counts at 1 kb bins) across the *C. albicans* genome. **(B)** Correlation between RNAPII occupancy (log10 counts) and transcriptional levels (RNA-seq; log10 counts) at protein-coding genes. **(C)** Histone H3 is depleted at centromeric regions. Fold enrichment (log2) of histone H3 relative to unmodified H4 across CEN1 (Chr1) and CEN5 (Chr5) centromeric and pericentromeric regions in *C. albicans*. The Cse4^CENP-A^ enrichment profile [43] is shown as comparison. The blue bar indicates statistically significant depleted regions for histone H3.

### H3K4me^3^, H3K9Ac and H4K16Ac mark C. albicans active genes

To delineate the chromatin signature of protein-coding *C. albicans* genes, enrichment profiles for each histone modification were compared to histone H4. Differential enrichment testing using DESeq2 allowed the identification of regions with statistically significant enrichment or depletion for particular histone marks compared to histone H4. We annotated these loci by proximity to annotated protein coding genes and non-coding RNAs [25–27]. For RNAPII, aligned reads from ChIP (IP) samples were normalised to aligned reads from the matching input (I) sample. Metagene analyses demonstrate that, as expected, RNAPII is enriched across all gene bodies while unmodified histone H3 is not significantly enriched or depleted relative to unmodified histone H4. In contrast, H3K4me^3^ and H3K9Ac are more prominent at the transcriptional start site (TSS) and 5’ regions of genes, and H4K16Ac is enriched at gene bodies. (Fig 2A).

**Figure 2.**
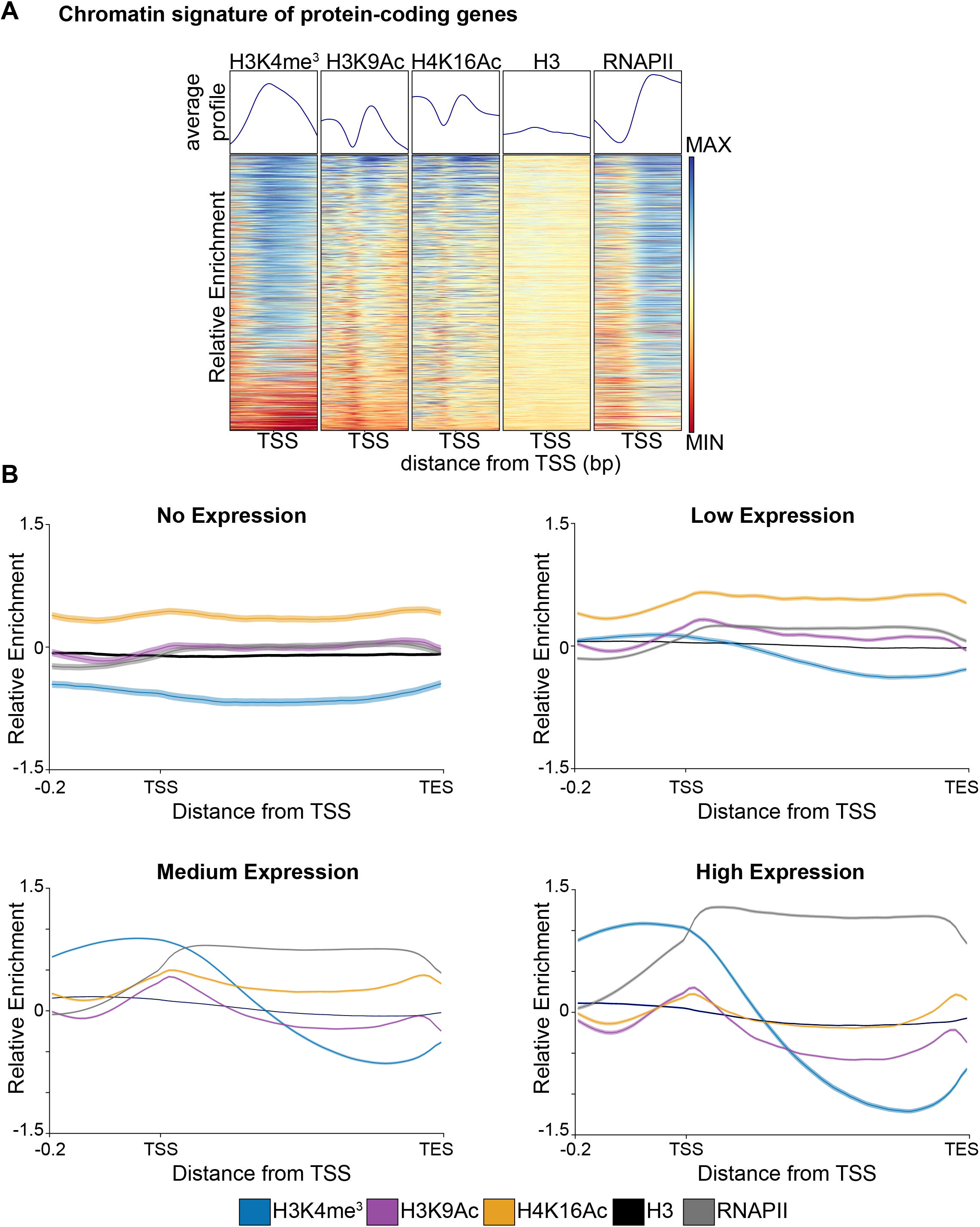
*C. albicans* chromatin modifications mirror their transcriptional state **(A)** Chromatin signature of *C. albicans* genes (n=6408). Average profiles and heatmaps of histone modification signatures around the Transcriptional Start Sites (TSS) of genes. The relative fold enrichment (log2) for each histone modification normalised to unmodified histone H4, or aligned reads of immunoprecipitated (IP) sample normalised to aligned reads of Input sample (for RNAPII ChIP-seq) is displayed within a region spanning ±0.5□kb around TSS. The gradient blue-to-red colour indicates high-to-low enrichment in the corresponding region. Min: - 1.5 log2. Max: +1.5 log2. **(B)** Average profiles of histone modifications and RNAPII occupancy across gene sets of different expression level (no [n = 416], low [n = 1369], medium [n = 3570] and high [n = 983] expression). For each histone modification, the fold enrichment (log2) relative to unmodified H4 is shown. For RNAPII the IP/I enrichment (log2) is shown.

To further explore the relationship between chromatin modifications and gene transcriptional states, we grouped all genes into four sets based on expression level (no expression, low expression, medium expression and high expression) as revealed by RNA-seq analysis (Fig S3). Enrichment profile plots of the levels of histone modifications for each of these gene sets demonstrated that H3K4me^3^, H3K9Ac and H4K16Ac levels are very low at genes with low transcription rates. Levels of all modifications increase with increased gene expression reaching a maximum at highly transcribed genes (Fig 2B).

Therefore, in *C. albicans*, H3K4me^3^, H3K9Ac and H4K16Ac correlate with gene transcription; H3K4me^3^ and H3K9Ac are more enriched at the 5’ of a gene and H4K16Ac at the gene bodies.

### γ-H2A is enriched at convergent genes and in proximity of DNA replication origins

Having established that different regions of the *C. albicans* genome are marked by different chromatin modifications depending on their transcriptional state (Fig 2), we sought to systematically map the genome-wide profile of γH2A (γ-sites) in cycling undamaged cells, as this is a useful method to identify recombination-prone unstable sites [14]. Genome-wide ChIP-seq of γH2A identified 168 γ-sites where γH2A is enriched compared to histone H4 (Dataset S1). *C. albicans γ*-sites are different from the γH2A -domain caused by irrecoverable DSBs as γ-sites have generally a single peak of enrichment and they are shorter (average length 850 bp) than the 50-kb length of the that γH2A domain surrounding DSBs [44].

Analysis of γ-sites indicates that they are present at three classes of genomic loci: *(i)* longer genes that are often convergent, *(ii)* origins of replication and *(iii)* subtelomeric regions (discussed below) (Fig 3A and 3B).

**Figure 3.**
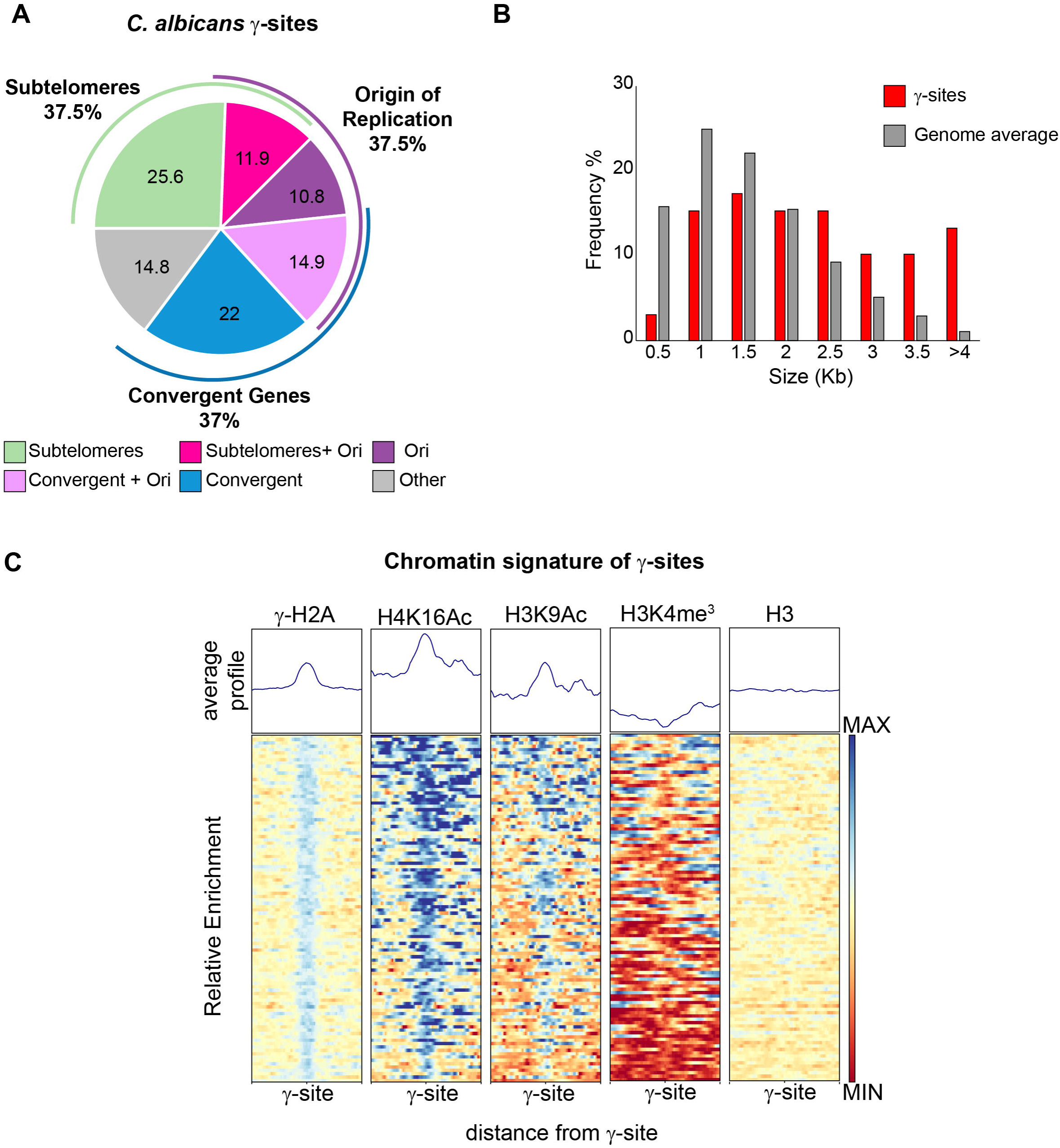
Identification of *C. albicans γ*-sites **(A)** Locations and frequencies of γ-sites throughout the *C. albicans* genome. **(B)** γ-sites map to longer genes. Histogram showing the gene lengths of γ-sites (red) compared to the genome average (grey) **(C)** Chromatin signature of γ-sites throughout the *C. albicans* genome. Average profiles and heatmaps of histone modification signatures at γ-sites. The relative fold enrichment (log2) for each histone modification normalised to unmodified histone H4 is displayed within a region spanning ±2□kb around the γH2A peak summits. The gradient blue-to-red colour indicates high-to-low enrichment in the corresponding region. Min: - 1.5 log2. Max: + 1.5 log2.

At convergent genes, γH2A enrichment is detected at both the gene bodies and intergenic regions, and no correlation was detected between gene expression levels and γH2A occupancy. Although we did not observe any correlation between γ-sites and histone H3 occupancy (Pearson Correlation r = 0.062), γ-sites are more likely to mark genomic regions that are acetylated on H4K16 and H3K9 (Pearson correlation r = 0.461 and 0.276, respectively). We also detect a weak negative correlation between γH2A occupancy and H3K4me^3^ (Fig 3C).

### The chromatin state of the C. albicans repetitive genome

Having determined the chromatin marks associated with *C. albicans* coding genes, we analysed the chromatin state of the *C. albicans* repetitive genome focusing on the major classes of DNA repeats: subtelomeric regions, the rDNA locus, MRS repeats and transposable elements (LTR and non-LTR retrotransposons). Sequence analysis of these elements can be problematic because of incomplete sequencing and their repetitive nature [25,29]. To estimate the chromatin modification state of these loci, we adopted a method previously applied to *S. cerevisiae* repeats and assumed that each repeat contributes equally to read-depth [45]. Consequently, reads that could not be uniquely mapped to one location were randomly assigned to copies of that repeat.

To investigate the chromatin state associated with the 16 subtelomeric regions in *C. albicans*, we analysed the ChIP-seq datasets in the 20-kb terminal regions of each chromosome arm. At these locations, occupancy of unmodified histone H3 was similar to histone H4 occupancy (Fig 4A, S4 and S5). In contrast, we detected large domains of chromatin that are hypomethylated on H3K4 and hypoacetylated on H3K9 and H4K16 (Fig 4A, S4 and S5). However, the H3K4 methylation and H3K9/H4K16 acetylation state of subtelomeres is not uniform as patches of high H3K4me^3^, H3K9Ac and H4K16Ac are detected within each subtelomere (Fig 4A, S4 and S5). We detected statistically significant γH2A enrichment at 13/16 subtelomeres (Fig 4A, S4 and S5). We suspect that absence of γ-sites at ChrRR, Chr1R and Chr7L subtelomeric regions is due to incomplete genome assembly [25,29]. Subtelomeric γH2A enrichment is not uniform but present at distinct peaks within each subtelomere, which largely associate with hypoacetylated and hypomethylated chromatin (Fig 4A, S4 and S5).

**Figure 4.**
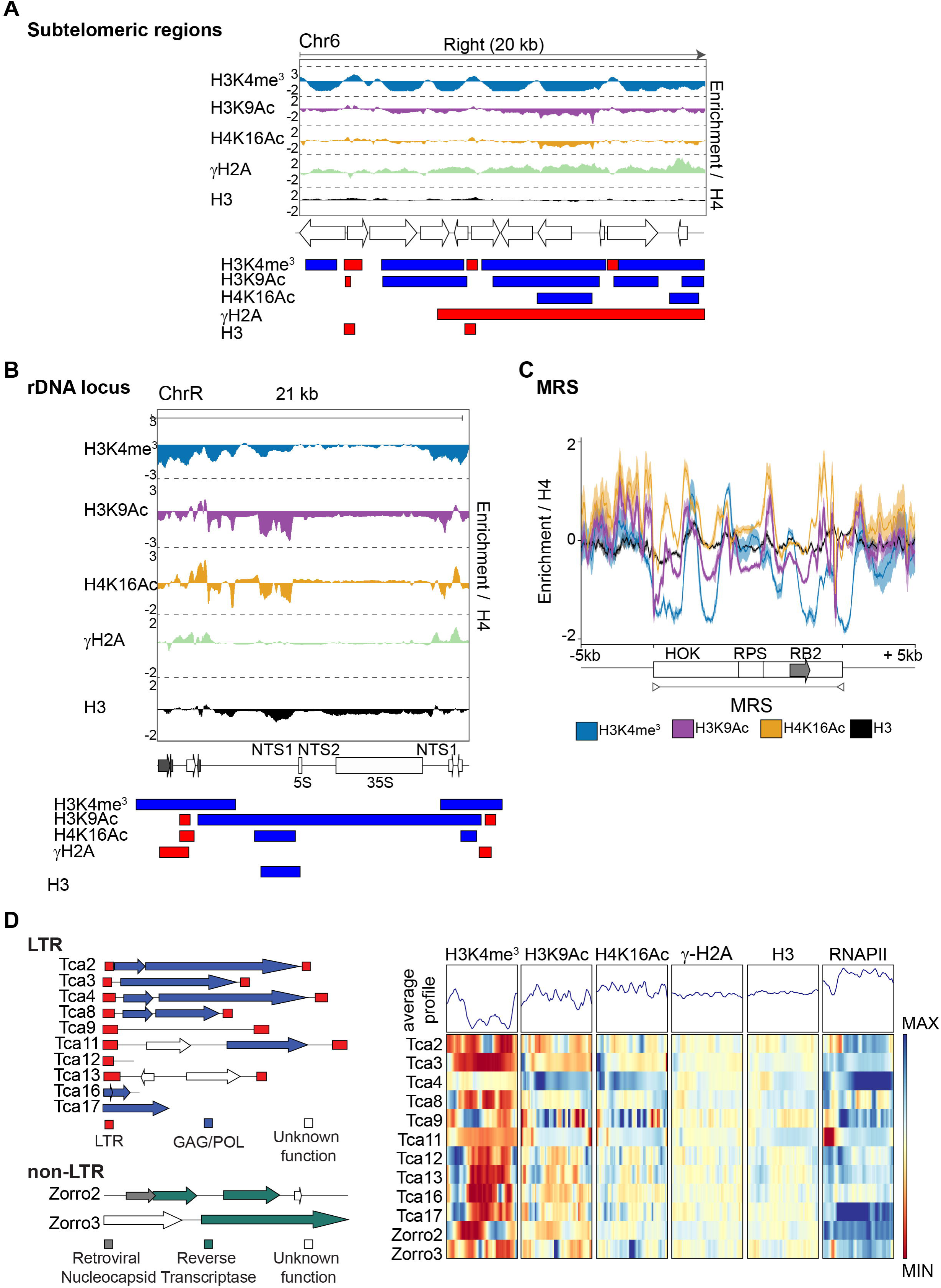
Chromatin signature of *C. albicans* repetitive elements **(A)** *Top*: Fold enrichment (log2) of H3K4me^3^ H3K9Ac, H4K16Ac, γH2A and H3 relative to unmodified H4 across the 20 kb right terminal region of chromosome 6 (Chr6). *Middle*: Diagram of coding genes found at these regions, according to assembly 22. *Bottom*: Diagram depicting statistically significant enriched (red) or depleted (blue) domains for each histone modification. **(B)** *Top*: Fold enrichment (log2) of H3K4me3, H3K9Ac, H4K16Ac, γH2A and H3 relative to unmodified H4 at the rDNA locus and flanking regions (ChrR). *Middle*: Diagram of coding genes (white) and ncRNA (grey) found at this region, according to assembly 22. *Bottom*: Diagram depicting statistically significant enriched (red) or depleted (blue) domains for each histone modification **(C)** Average profiles of histone modifications at MRS repeats, and upstream and downstream sequences. The grey arrow indicates the location of the FGR gene. For each histone modification, the fold enrichment (log2) relative to unmodified H4 is shown. **(D)** *Left:* Diagrams of the structure of the *C. albicans* LTR and non-LTR retrotransposons. *Right:* Chromatin signature of LTR and non-LTR retrotransposons. Average profiles and heatmaps of histone modification signatures across each sequence. The relative fold enrichment (log2) for each histone modification normalised to unmodified histone H4, or aligned reads of immunoprecipitated (IP) sample normalised to aligned reads of Input sample (for RNAPII ChIP-seq) is displayed. The gradient blue-to-red colour indicates high-to-low enrichment in the corresponding region. Min: - 1.5 log2. Max: + 1.5 log2.

Analysis of chromatin modifications associated with the rDNA locus demonstrate that the NTS1 and NTS2 regions are assembled into a chromatin structure resembling heterochromatin where nucleosomes are hypomethylated on H3K4 and hypoacetylated on H3K9 and H4K16 (Fig 4B). These findings are consistent with our published results demonstrating that these regions are assembled into transcriptionally silent heterochromatin [46]. Intriguingly, we detected two γ-sites at convergently transcribed genes surrounding the rDNA locus (Fig 4B).

This analysis also reveals that MRS repeats and retrotransposons (LTR and non-LTR) are associated with chromatin that is largely hypomethylated on H3K4 (Fig 4C, 4D). In contrast, H3K9Ac and H4K16Ac are similar to histone H4 levels (Fig 4C, 4D). We did not detect any statistically significant enrichment of γ-H2A at either MRSs or retrotransposons.

Therefore, different *C. albicans* repetitive elements are associated with distinct chromatin states. Repetitive regions are more likely to be hypomethylated on H3K4, but are neither hypoacetylated on H3K9 and H4K16 nor enriched for γ-H2A.

### The HDAC Sir2 governs the hypoacetylated state associated with C. albicans rDNA locus and subtelomeric regions

We have previously shown that, in *C. albicans*, the histone deacetylase Sir2 maintains the low level of H3K9Ac associated with the NTS regions of the rDNA locus [46]. To assess the role of the HDAC Sir2 in maintaining acetylation levels across the *C. albicans* genome, we performed H3K9Ac and H4K16Ac ChIP-seq analyses in WT and *sir2Δ/Δ* strains.

Traditional ChIP-seq are not inherently quantitative as it allows comparison of protein occupancies at different positions within a genome but it does not allow direct comparisons between samples derived from different strains [47–49]. To overcome this issue, we adapted *C. albicans* to a quantitative ChIP-seq (q-ChIP-seq) methodology [47–49]. To this end, WT and *sir2Δ/Δ* were spiked-in, at the time of fixation, with a single calibration sample from *S. cerevisiae* (Fig 5A). *S. cerevisiae* genome is a desirable exogenous reference for *C. albicans* cells because its genome is well studied and has a high-quality sequence assembly [50]. Moreover, reads originating from *C. albicans* or *S. cerevisiae* can be easily separated at the analysis level and our experiments reveal less than 2% of the total number of reads cannot be uniquely mapped (Table S2). Finally, histone proteins are well conserved between *C. albicans* and *S. cerevisiae* (Fig S1) and therefore the same histone antibody is likely to immunoprecipitate *C. albicans* and *S. cerevisiae* chromatin with the same efficiency.

**Figure 5.**
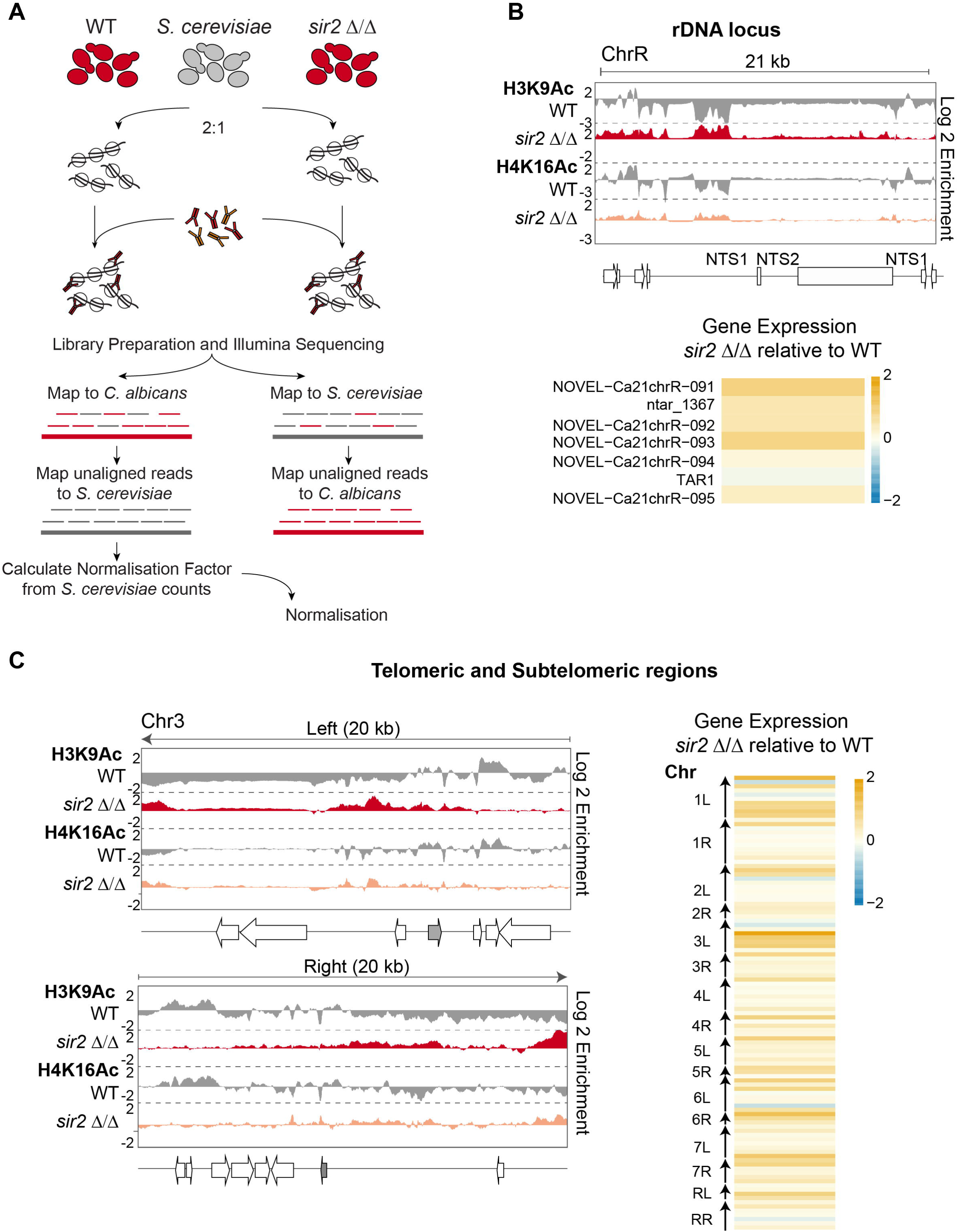
The HDAC Sir2 controls the chromatin state of subtelomeres and the rDNA locus **(A)** Schematic of the quantitative ChIP-seq experimental and analytical workflow. **(B)** *Top*: Fold enrichment (log2) of H3K9Ac and H4K16Ac relative to unmodified H4 in WT cells, and relative to WT in *sir2Δ/Δ* cells, across the rDNA loci of chromosome R (ChrR). *Middle*: Diagram of transcripts found at this region, according to assembly 22. *Bottom*: Heatmap depicting changes in gene and ncRNA expression across the rDNA region in *sir2Δ/Δ* cells relative to WT. The gradient yellow-to-blue colour indicates high-to-low expression. Min: - 2 log2. Max: + 2 log2. **(C)** *Left*: Fold enrichment (log2) of H3K9Ac and H4K16Ac relative to unmodified H4 in WT cells, and relative to WT in *sir2Δ/Δ* cells, across the 20 kb left and right terminal regions of chromosome 3 (Chr3). Diagrams of coding genes (TLO: grey) found at these regions, according to assembly 22, are below. *Right*: Heatmap depicting changes in gene and ncRNA expression in *sir2Δ/Δ* cells relative to WT at the 10 kb terminal regions of all *C. albicans* chromosomes. The gradient yellow-to-blue colour indicates high-to-low expression. Min: - 2 log2. Max: + 2 log2.

The q-ChIP-seq analyses identify only two regions of the *C. albicans* genome with increased H3K9Ac and H4K16Ac levels: subtelomeric regions and the NTS region of the rDNA locus (Fig 5B, 5C, S6, S7 and Dataset S1). Deletion of *SIR2* does not lead to increased histone acetylation levels at euchromatic regions or at other repetitive elements such as MRS and retrotransposons (Dataset S1). In agreement with these findings, the majority (83%) of gene expression changes observed in *sir2Δ/Δ* cells occur at the rDNA locus and subtelomeric regions (Fig 5B, 5C, Dataset S1 and [46]) We conclude that the *C. albicans* HDAC Sir2 acts exclusively at two genomic regions: the rDNA locus and subtelomeric regions. Our findings are consistent with the hypothesis that Sir2-mediated histone deacetylation represses gene expression at these locations.

### Set1-dependent methylation of H3K4 impacts gene expression differentially at different repeats

Our data demonstrates that *C. albicans* repetitive elements are associated with chromatin that is hypomethylated on H3K4. However, at these regions, H3K4 methylation is not completely ablated as pockets of H3K4me^3^ are detected (Fig 4). In *S. cerevisiae* and *S. pombe*, the H3K4 methyltransferase Set1 has been implicated in both gene repression and activation [51–56]. *S. cerevisiae* Set1 also maintains the transcriptional silencing associated with heterochromatic regions such as the telomeres and the rDNA locus [51–56]. *C. albicans* Set1 is important for efficient yeast-to-hyphae switching but its function in regulating chromatin structure and gene expression is unknown [57].

To gain insights into the role of *C. albicans* Set1, we performed H3K4me^3^ q-ChIP-seq and RNA-seq analyses of WT and *set1Δ/Δ* strains. *C. albicans* Set1 clearly plays a major role in maintaining chromatin structure as 6846 loci, scattered throughout the genome, have a statistically significant reduction of H3K4me^3^ in *set1Δ/Δ* compared to WT strain (Fig 6A and Dataset S1). RNA-seq analysis reveals that Set1 regulates gene expression both positively and negatively, as genes with a reduced H3K4me^3^ pattern can be either upregulated (2320 genes/ ncRNAs) or downregulated (3184 genes/ ncRNAs) in *set1Δ/Δ* compared to WT (Fig 6A and Dataset S1). Analyses of the H3K4me^3^ pattern and gene expression levels associated with repetitive elements demonstrates that Set1 has distinct roles at different repeats. At subtelomeric regions and the rDNA locus, deletion of the *SET1* gene leads to the reduced H3K4me^3^ levels and is accompanied by the down-regulation of associated genes (Fig 6B, 6C and S8). In contrast, the reduced H3K4me^3^ pattern associated with MRS repeats in the *set1Δ/Δ* strain leads to increased expression of coding and non-coding RNAs originating from MRS repeats (Fig 6D). Finally, deletion of *SET1* leads to decreased H3K4me^3^ at retrotransposons without a significant impact on expression of retrotransposon-associated coding and non-coding RNAs.

**Figure 6.**
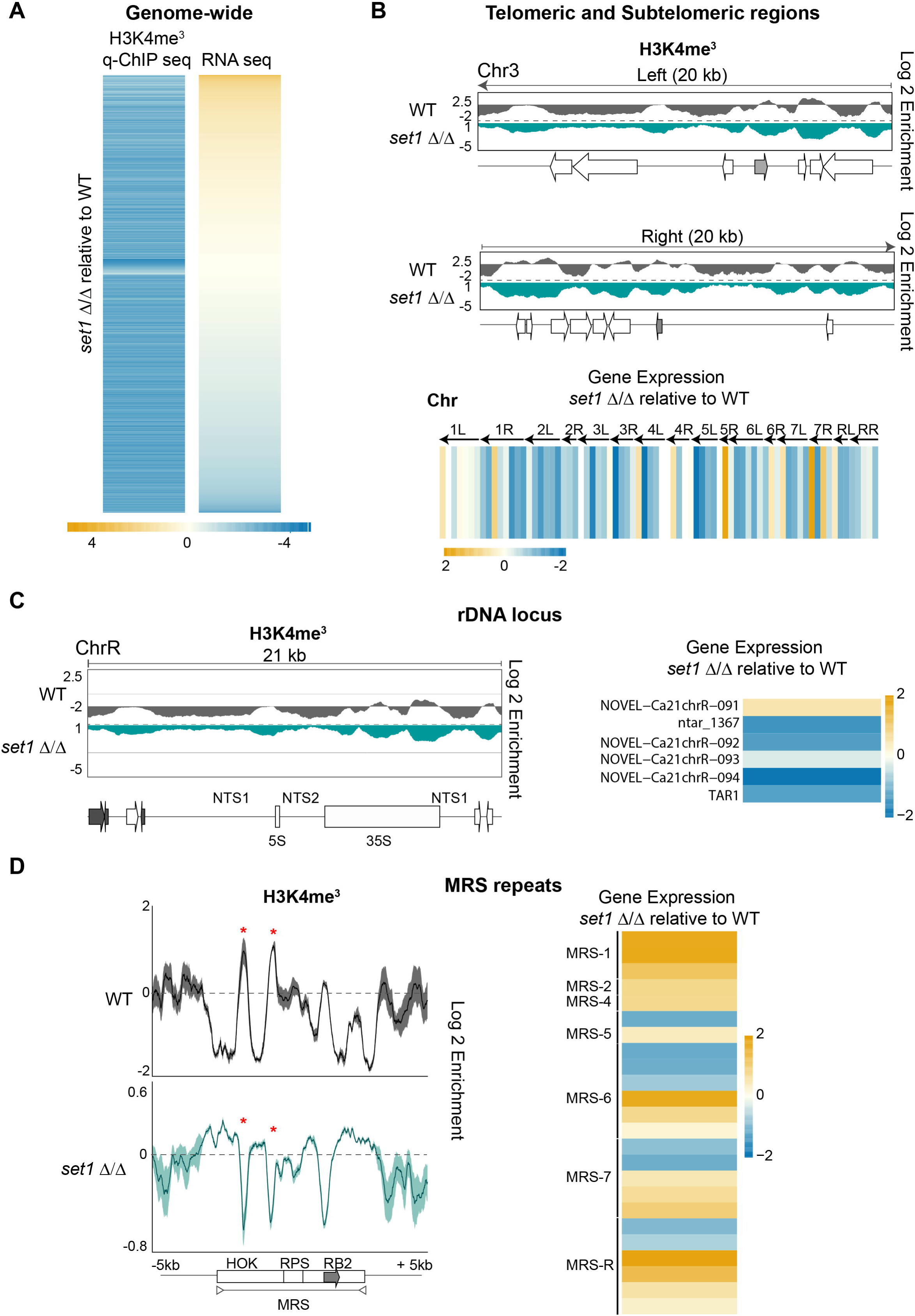
Chromatin and gene expression changes of set1Δ/Δ strain **(A)** Heatmap depicting changes in expression of genes and ncRNA associated with statistically significant changes in H3K4me^3^ enrichment in *set1Δ/Δ* cells relative to WT cells. The gradient yellow-to-blue colour indicates high-to-low enrichment/expression. Min: - 4 log2. Max: + 4 log2 **(B)** *Top*: Fold enrichment (log2) of H3K4me^3^ relative to unmodified H4 in WT cells, and relative to WT in *set1Δ/Δ* cells, across the 20 kb left and right terminal regions of chromosome 3 (Chr3). Diagrams of coding genes (TLO: grey) found at these regions, according to assembly 22, are below. *Bottom*: Heatmap depicting changes in gene and ncRNA expression in *set1Δ/Δ* cells relative to WT at the 10 kb terminal regions of all *C. albicans* chromosomes. The gradient yellow-to-blue colour indicates high-to-low expression. Min: - 2 log2. Max: + 2 log2. **(C)** *Left*: Fold enrichment (log2) of H3K4me^3^ relative to unmodified H4 in WT cells, and relative to WT in *set1Δ/Δ* cells, across the rDNA loci of chromosome R (ChrR). Diagrams of coding genes and ncRNA (grey) found at this region, according to assembly 22, are below. *Right*: Heatmap depicting changes in gene and ncRNA expression across the rDNA region in *set1Δ/Δ* cells relative to WT. The gradient yellow-to-blue colour indicates high-to-low expression. Min: - 2 log2. Max: + 2 log2. **(D)** *Left*: Profiles of fold enrichment (log2) of H3K4me^3^ relative to unmodified H4 in WT cells, and relative to WT in *set1Δ/Δ* cells averaged across the MRS repeats, and the upstream and downstream sequences. The grey arrow indicates the location of the FGR gene. *Right*: Heatmap depicting changes in gene and ncRNA expression across all of the MRS regions in *set1Δ/Δ* cells relative to WT. The gradient yellow-to-blue colour indicates high-to-low expression. Min: - 2 log2. Max: + 2 log2.

We conclude that Set1 is the major H3K4-methyltransferase *in C. albicans* playing a key role in controlling chromatin structure and gene expression. Importantly, our analysis reveals that although deletion of *SET1* results in decreased H3K4 methylation across all repetitive elements, Set1 influences gene expression differentially at each repetitive element.

## DISCUSSION

Here, we present the first comprehensive chromatin profiling of histone modifications associated with the *C. albicans* genome. Furthermore, we present the first *C. albicans* quantitative ChIP-seq to delineate the role of the chromatin modifiers Sir2 and Set1 in *C. albicans*.

### The chromatin state of the C. albicans repetitive and non-repetitive genome

In all organisms, gene-rich genomic regions are associated with a histone modification pattern mirroring their transcriptional state where H3K4me^3^ and H3K9Ac are enriched at active gene promoters and H4K16Ac is localised at gene bodies of expressed genes [9,10]. Our first objective was to obtain “proof of concept” epigenomic maps of chromatin modifications associated with gene rich regions of the *C. albicans* genome. A robust histone modification profiling relies on *(i)* the use of antibodies that recognise modified histones with high specificity and *(ii)* the use of appropriate biological controls. Specificity of antibodies used in this study has been tested in *S. pombe* or *S. cerevisiae* histone mutants lacking the modifiable amino acid (H3K9, H4K16, H2AS129) [14,58]. To distinguish between nucleosome occupancy and depletion/enrichment of specific histone modifications ChIP seq analyses was also performed using antibodies recognising unmodified histone H3 and H4. This is an important control that should be included in all studies aimed to analyse chromatin modification genome wide.

Our results confirm the validity of our experimental approach and conform to the chromatin pattern reported in other organisms by showing that active genes, associated with high levels of RNA Pol II, are assembled into canonical euchromatin where H3K4me^3^ and H3K9Ac are associated with promoters and H4K16Ac is enriched at gene bodies. We conclude that in *C. albicans*, as in other organisms, a specific histone modification pattern is predictive of active transcription.

Analyses of the chromatin profiling of repetitive elements demonstrate that repetitive regions associated with the NTS regions of the rDNA locus and subtelomeric regions are packaged into chromatin resembling the heterochromatic structure of other organisms, like the budding yeast *S. cerevisiae*, lacking H3K9me/H3K27me systems [45]. These findings are in agreement with our previous study demonstrating that the rDNA locus and subtelomeric regions are able to silence embedded marker genes, which is a hallmark of heterochromatic regions [46]. In contrast, we find that *C. albicans* retrotransposons and MRS repeats are assembled into a distinct chromatin state where nucleosomes are hypomethylated on H3K4me^3^, but also acetylated on H3K9 and H4K16. Analyses of clinical isolates and *in vivo* evolution experiments have demonstrated that, in the host, MRSs and transposons are recombination hotspots as they are known sites of translocations [32,38,59]. Given the key roles of chromatin in regulating genome stability, it will be important to investigate whether the chromatin packaging of MRS and transposons regulates genome stability.

### γH2A decorates C. albicans heterochromatic regions and potential recombination-prone unstable sites

The genome-wide γ-H2A profiling performed in this study reveals that this histone modification, a hallmark of DNA damage, is enriched at heterochromatic regions assembled into hypoacetylated chromatin that is also hypomethylated on H3K4. This is similar to observations in other fungal organisms where γ-H2A decorates heterochromatic regions [14–17]. In contrast, we did not detect any significant enrichment of γ-H2A at other repetitive elements such as MRS repeats and transposable elements. This is surprising because, in the host, MRS repeats are recombination hot-spots [32,38] and therefore a place where DSBs might be expected to accumulate. *C. albicans* genome instability is increased under host-relevant stresses [60,61] and therefore we propose that the recombination potential of MRSs is unlocked following exposure the specific host niche stresses.

Finally, we detected 168 additional γ-sites located in proximity of origins of replication or convergent genes that are often long. DNA replication origins are known replication fork barriers in many organisms and read-through transcription of convergent genes can also cause genome instability by, for example, R-loop formation [62]. Therefore, we propose that the γ-sites identified in this study represent novel recombination-prone unstable sites of the *C. albicans* genome.

### The role of the histone deacetylase Sir2 and the histone methyltransferase Set1 in controlling the C. albicans epigenome

We present the first quantitative ChIP-seq in *C. albicans* that has allowed us to delineate the roles of the histone modifying enzymes Sir2 and Set1. We demonstrate that Sir2 maintains the hypoacetylated state of heterochromatic regions associated with the rDNA locus and subtelomeric regions. Sir2 deacetylation at these loci is linked to gene repression as shown by RNA-seq analysis. In contrast, we find that deletion of Sir2 does not lead to increased histone acetylation and gene expression at other genomic regions. Two possible scenarios could explain these findings: *(i*) Sir2 is specifically targeted to subtelomeres and the rDNA locus or *(ii)* other histone deacetylases act redundantly to Sir2 regulating hypoacetylation and gene expression at other genomic locations.

We present evidence demonstrating that the HMT Set1 is a major contributor to chromatin structure in *C. albicans*. Indeed, deletion of *SET1* leads to an almost complete ablation of H3K4 methylation and is linked to extensive gene expression changes. This demonstrates that *C. albicans* Set1 is the major H3K4 methyltransferase. It is particularly intriguing that deletion of *SET1* leads to decreased H3K4me^3^ at all known repeats, yet its effect on gene expression can be the opposite. Indeed, we demonstrate that at the rDNA locus and subtelomeric regions Set1 represses gene expression while it activates gene expression at MRS repeats. Further studies will untangle the role of Set1 at different genomic regions.

## CONCLUSIONS

In this study we present the first epigenomic map of histone modifications associated with the *C. albicans* genome. Given the key role of chromatin in regulating *C. albicans* biology, the data generated in this study provide an invaluable resource to a better understanding of this important human fungal pathogen.

## METHODS

### Yeast growth and manipulation

Strains used in this study are listed in the Table S1. Yeast cells were cultured in YPAD broth containing 1% yeast extract, 2% peptone, 2% dextrose, 0.1 mg/ml adenine and 0.08 mg/ml uridine at 30°C.

### Antibody Information

The following antibodies were used in this study: anti-H2AS129p (Millipore; Cat No: 07-745-I), anti-H3 (Abcam; Cat No: ab1791), anti-H4 (Millipore; Cat No: 05-858), anti-H3K4me3 (Active Motif; Cat No: 39159), anti-H3K9ac (Active Motif; Cat No: 39137), anti-H4K16ac (Active Motif; Cat No: 39167), and anti-RNA Polymerase II (BioLegend; Cat No: 664903).

### ChIP-seq

Chromatin immunoprecipitation with deep-sequencing (ChIP-seq) was performed as follows: 5 ml of an overnight culture grown in YPAD was diluted into fresh YPAD and grown until the exponential phase (OD_600_ = 0.6-0.8). 20 OD_600_ units of cells were fixed with 1% formaldehyde (Sigma) for 15 minutes at room temperature. Reactions were quenched by the addition of glycine to a final concentration of 125 mM. Cells were lysed using acid-washed glass beads (Sigma) and a DisruptorGenie (Scientific Industries) for four cycles of 30 minutes at 4°C with 5 minutes on ice between cycles. Chromatin was sheared to 200–500 bp using a BioRuptor sonicator (Diagenode) for a total of 20 minutes (30 seconds on, 30 seconds off cycle) at 4°C.

Immunoprecipitation was performed overnight at 4°C using 2 µl of the appropriate antibody and 25 µl of protein G magnetic Dynabeads (Invitrogen). ChIP DNA was eluted, and cross-links reversed at 65°C in the presence of 1% SDS. All samples were then treated with RNaseA and proteinase K before being purified by phenol:chloroform extraction and ethanol precipitation. Libraries were prepared and sequenced as 50bp single end reads on an Illumina Hi seq2000 platform by the Genomics Core Facility at EMBL (Heidelberg, Germany). All ChIP-seq experiments were carried out in biological duplicates.

### q-ChIP-seq

Quantitative chromatin immunoprecipitation with deep-sequencing (q-ChIP-seq) was performed similarly to the ChIP-seq method, except: 5 ml of an overnight culture of the *S. cerevisiae* reference strain BY4741 was grown alongside *C. albicans* in YPAD. These cultures were then diluted into fresh YPAD and grown until the exponential phase (OD_600_ = 0.6-0.8). 20 OD_600_ units of *C. albicans* cells were combined with 10 OD_600_ units of *S. cerevisiae* cells, and then fixed with 1% formaldehyde (Sigma) for 15 minutes at room temperature. After the cells had been fixed, the q-ChIP-seq sample was processed as a single ChIP-seq sample throughout the experiment until completion of DNA sequencing. All q-ChIP-seq experiments were carried out in biological duplicates.

### RNA-seq

RNA was extracted from exponential cultures (OD_600_ = 0.6-0.8) using a yeast RNA extraction kit (E.Z.N.A. Isolation Kit RNA Yeast; Omega Bio-Tek) following the manufacturer’s instructions. RNA quality was checked by electrophoresis under denaturing conditions in 1% agarose, 1x HEPES, 6% formaldehyde (Sigma). RNA concentration was measured using a NanoDrop ND-1000 spectrophotometer. Strand-specific cDNA Illumina barcoded libraries were generated from 1 µg of total RNA and sequenced as 50bp single end reads using an Illumina Hi seq2000 sequencer by the Genomics Core Facility at EMBL (Heidelberg, Germany). All RNA-seq experiments were carried out in biological duplicates.

### Analysis of high-throughput sequencing

All datasets generated and analysed during the current study are available in the BioProject NCBI repository (https://www.ncbi.nlm.nih.gov/bioproject) under the BioProject ID (PRJNA503946).

#### ChIP-seq Analysis

Illumina reads were mapped using Bowtie2 [63] to a custom haploid version of assembly 22 of the *C. albicans* genome (Table S2). Reads that mapped to repeated sequences were randomly assigned to copies of that repeat, allowing for an estimation of enrichment at the repetitive elements of the genome. Peak calling was performed using MACS2 [64] on the default settings, except that no model was used with all reads extended to 250 bp. MACS2 was run separately on both biological replicates for each ChIP-seq sample. For each sample analysed with MACS2, the IP sample was the “treatment,” and the input sample was the “control.” We defined peaks as reproducible if they were called in both data sets. Read counts within peak intervals were generated using featureCounts [65]. For each interval, biological duplicate counts were compared between each histone modification and unmodified histone H4 samples using DESeq2, with an adjusted p-value threshold of <0.05 being used to identify significant differences. Replicates were compared by generating a raw alignment coverage track and performing a Pearson correlation between them using the multiBamSummary and plotCorrelation tools as part of the deepTools2 package (Fig S9) [66]. Genome coverage tracks were made using the pileup function of MACS2 [64] and tracks from biological replicates were averaged after the replicates were deemed to be sufficiently correlative (r > 0.9). For each coverage track, reads per million (RPM) were calculated. The histone modification coverage tracks were normalised to unmodified histone H4, and the RNAPII track was normalised to the respective input sample. All coverage tracks were visualised using IGV [67]. Metaplots and heatmaps were made using computeMatrix, plotProfile and plotHeatmap tools as part of the deepTools2 package [66].

#### q-ChIP-seq Analysis

To isolate the reads that uniquely aligned to the *C. albicans* genome, the full datasets were first aligned to the *S. cerevisiae* genome (sacCer3). The unaligned reads were output as separate fastq files, and then these reads were aligned to a custom haploid version of assembly 22 of the *C. albicans* genome (Table S2). The same strategy was used to isolate reads that uniquely aligned to *S. cerevisiae* (Table S2). All alignments were performed using Bowtie2 [63]. The unique *S. cerevisiae* reads were then used to calculate the normalisation factor (normalisation factor = 1 / [unique reference reads/1,000,000]), according to Orlando et al. [48]. Reads that mapped to repeated sequences in the *C. albicans* genome were randomly assigned to copies of that repeat. Peak calling was performed using MACS2 [64] on the default settings, except that no model was used with all reads extended to 250bp. MACS2 was run separately on both biological replicates for each ChIP-seq sample. For each sample analysed with MACS2, the IP sample was the “treatment,” and the input sample was the “control.” Peaks called in both replicate datasets for mutant and WT samples were combined into one peak set for each histone modification. Read counts within these peak intervals were generated using featureCounts (Liao et al. 2014), which were then scaled by the normalisation factor to obtain the reference reads per million (RRPM). For each interval, RRPM values were compared between the mutant and WT samples using a two-sample t-test, with a p-value threshold of <0.05 being used to identify significant differences. Replicates were compared by generating a raw alignment coverage track and performing a Pearson correlation between them using the multiBamSummary and plotCorrelation tools as part of the deepTools2 package (Fig S9) [66]. Genome coverage tracks were made using the pileup function of MACS2 [64] and for each track, RRPM values were calculated using the normalisation factor. Coverage tracks from biological replicates were averaged after the replicates were deemed to be sufficiently correlative (r > 0.9), and the mutant strain coverage tracks were normalised to the WT coverage. All tracks were visualised using IGV [67]. Metaplots and heatmaps were made using computeMatrix, plotProfile and plotHeatmap tools as part of the deepTools2 package [66].

#### RNA-seq Analysis

Reads were aligned to a custom haploid version of assembly 22 of the *C. albicans* genome using HISAT2 (Table SX) [68], and per-gene transcript quantification was performed using featureCounts, which discards multi-mapped read fragments; therefore, only uniquely mapped reads were included for the expression analysis [65]. Differential expression testing was performed using DESeq2, with an adjusted p-value threshold of <0.05 being used to determine statistical significance. Replicates were compared by generating a raw alignment coverage track and performing a Pearson correlation between them using the multiBamSummary and plotCorrelation tools as part of the deepTools2 package (Fig S10) [66]. Scatterplots and correlation analyses were performed in R using Pearson correlation.

## Supporting information

## DECLARATION

### Ethics approval and consent to participate

‘Not applicable’;

### Consent for publication’

Not applicable’;

### Availability of data and material

The datasets generated and analysed during the current study are available in the BioProject NCBI repository (https://www.ncbi.nlm.nih.gov/bioproject) under the BioProject ID (PRJNA503946).;

### Authors’ contributions

R.J.P conducted the ChIP seq, the RNA-seq and the bioinformatics analyses. E.W performed RNAseq experiments of *sir2 Δ/Δ* strain. A.B and J.B conceived the project, designed the experiments and wrote the manuscript.

## Acknowledgements

We thank members of the Kent Fungal Group, Jan Soetaert and Alison Pidoux for discussion and critical reading of the manuscript. We thank the Gene Core Facility at EMBL (Heidelberg-Germany) for Illumina Sequencing.

## ADDITIONAL FILES

Additional file 1-10: Fig S1 to S10. pdf

Additional file 11: Table S1: Strains used in this study

Additional file 12: Table S2.xsl; Sequencing and coverage information

Additional file 13: Dataset S1.xsl; Datasets generated in this study

