## Supplementary material for "Chromatin profiling of the repetitive and non-repetitive genome of the human fungal pathogen Candida albicans"

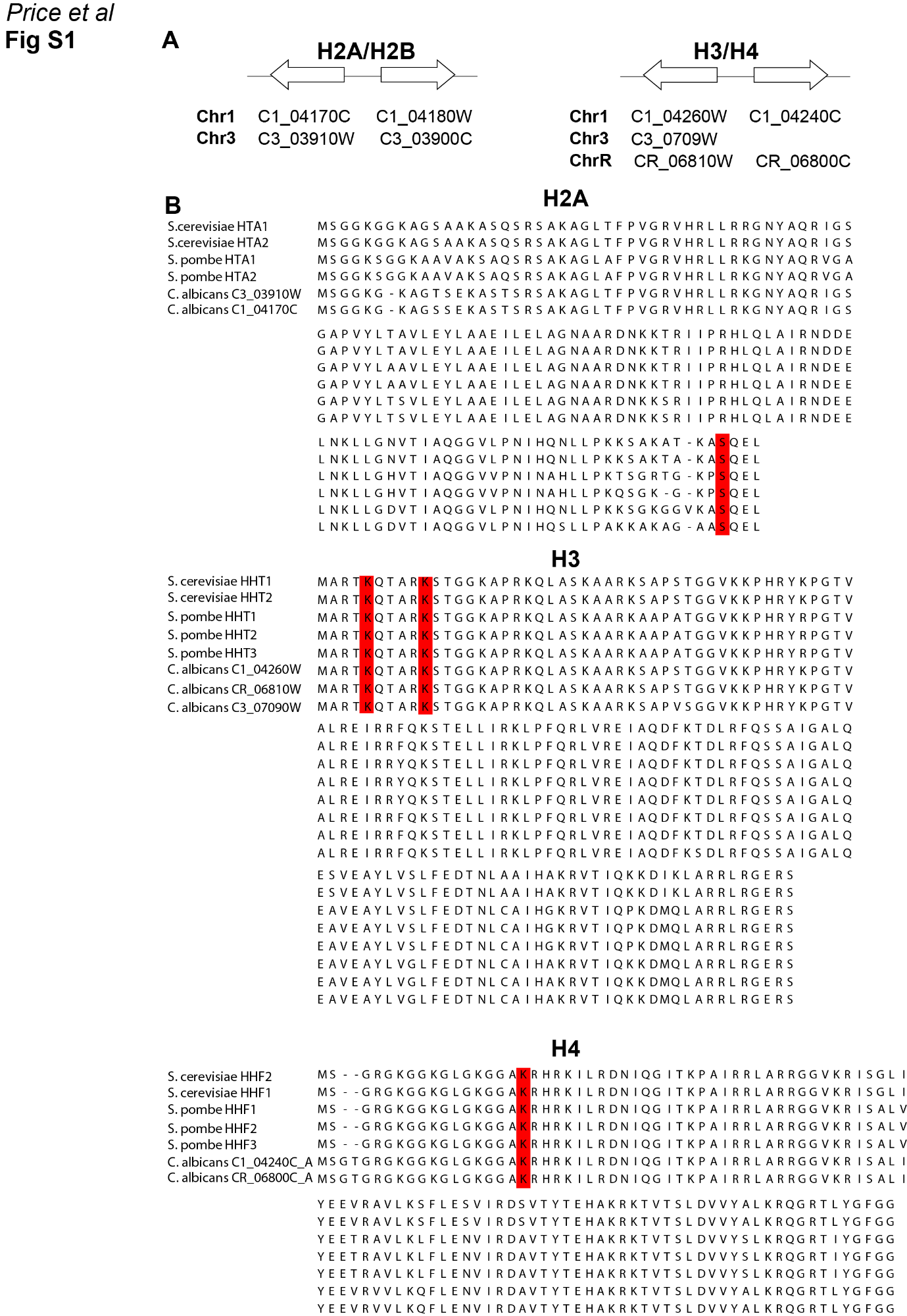

**Fig S1 (A)** *Top:* Diagram of genome organisation of *C. albicans* histone genes. *Bottom:* Histone gene names and locations, according to assembly 22. **(B)** Alignment of *C. albicans* H2A, H3 and H4 protein sequences against *S. cerevisiae* and *S. pombe* homologues. Residues highlighted in red are investigated in this study and are conserved across all species.

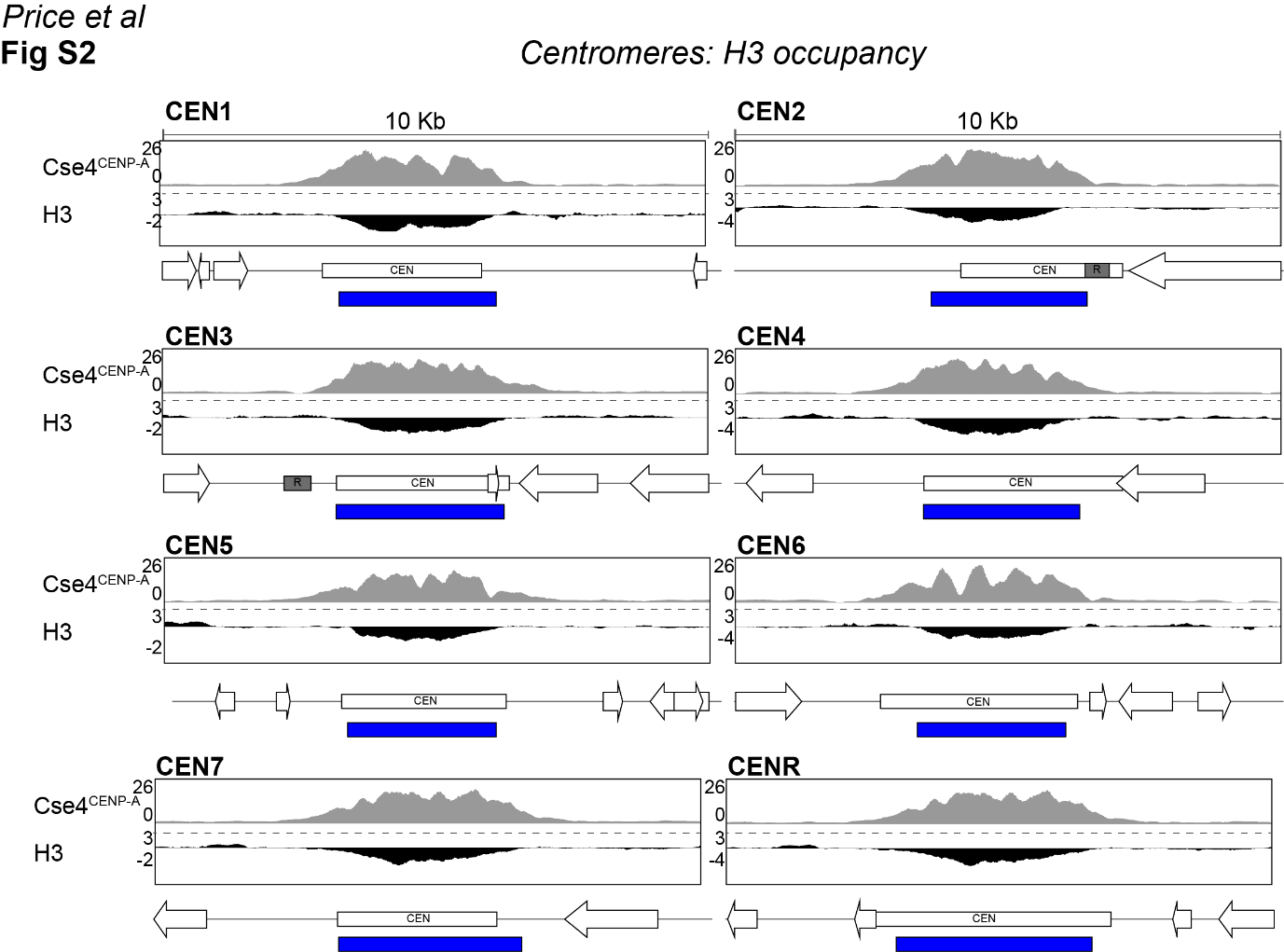

**Fig S2** Fold enrichment (log2) of histone H3 relative to unmodified H4 across the centromeric and pericentromeric regions of all 8 chromosomes in *C. albicans*. The CenpA enrichment profile is shown as comparison. The blue bars indicate statistically significant depleted regions for histone H3.

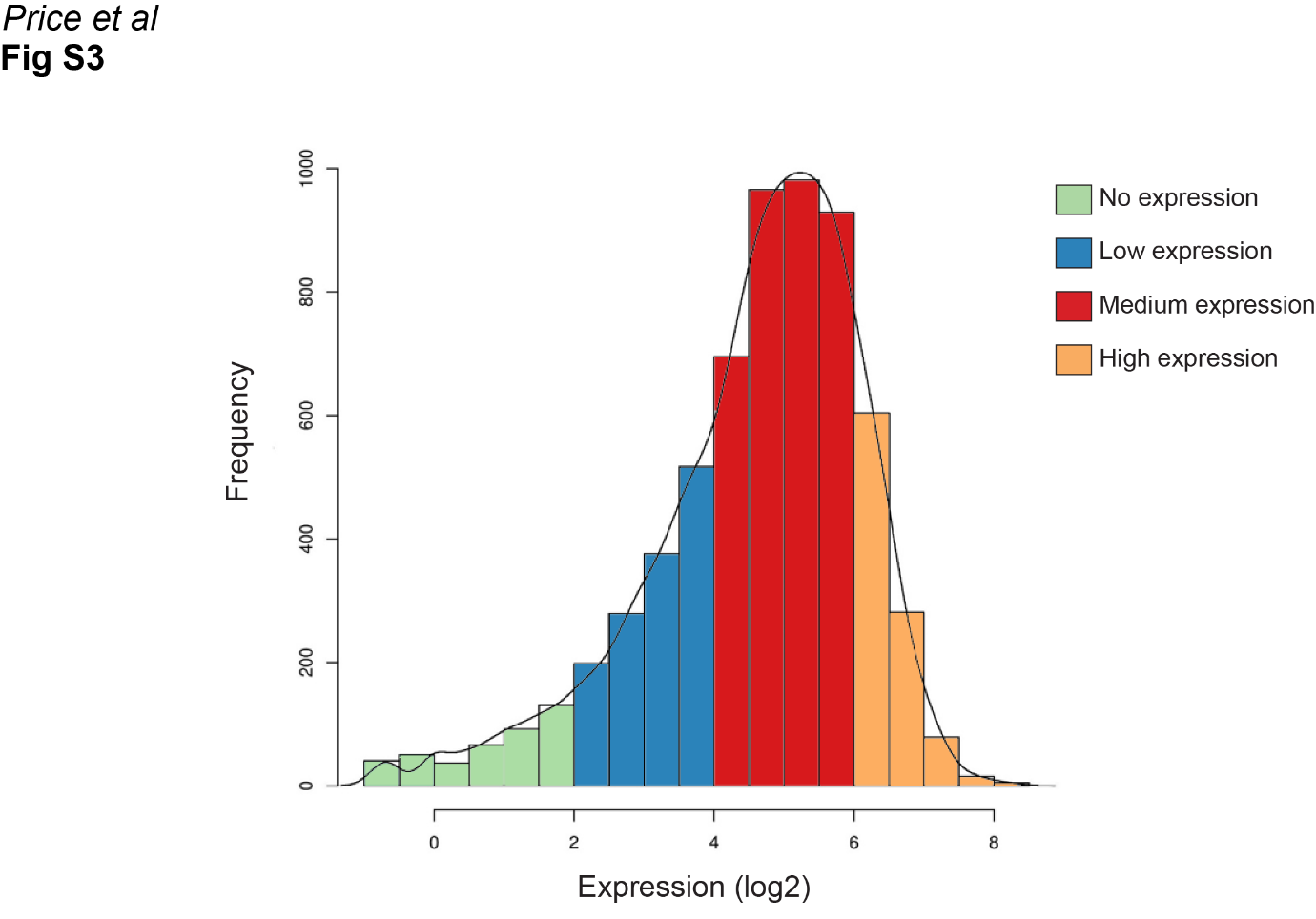

**Fig S3** Histogram of RNA-seq counts (log2) in WT cells across all protein coding genes, according to assembly 22. Genes were grouped into four sets organised by expression level; no expression (green, log2 counts <2, n = 416), low expression (blue, log2 counts 2-4, n = 1369), medium expression (red, log2 counts 4-6, n = 3570) and high expression (yellow, log2 counts >6, n = 983).

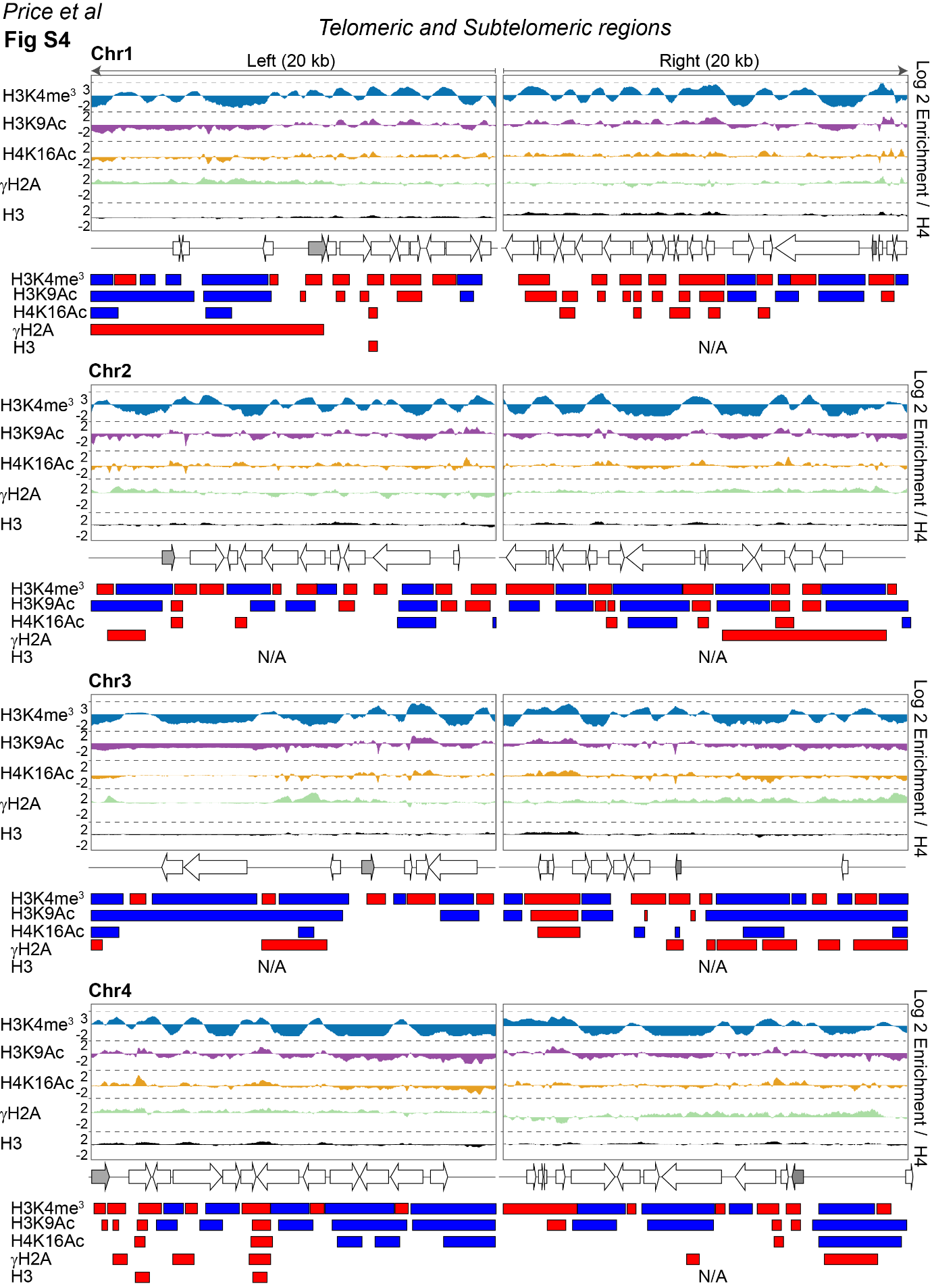

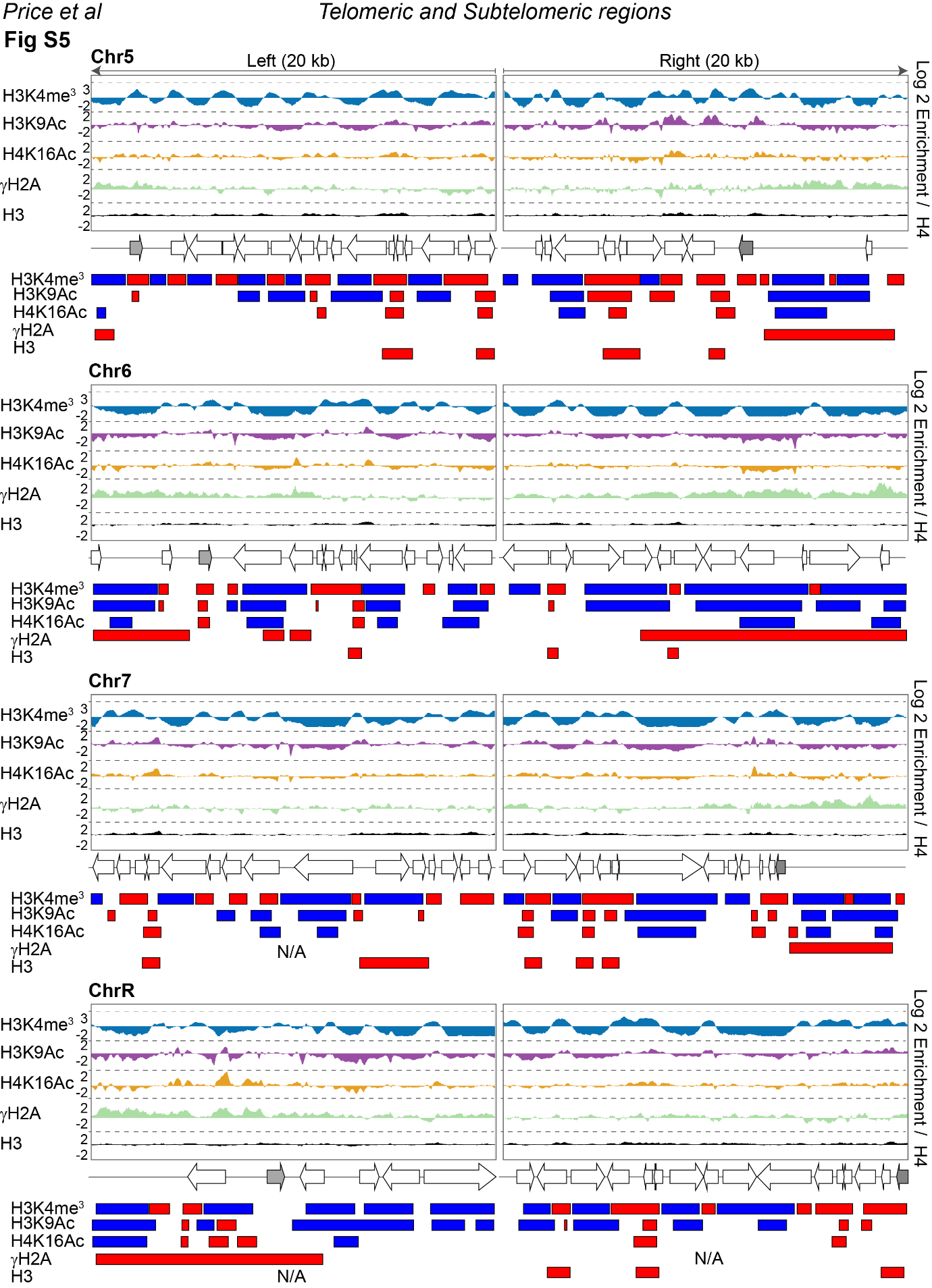

**Fig S4** and **Fig S5** *Top*: Fold enrichment (log2) of H3K4me^3^_,_ H3K9Ac, H4K16Ac, γH2A and H3 relative to unmodified H4 across the 20 kb left and right terminal region of each of the 8 *C. albicans* chromosomes. *Middle*: Diagrams of coding genes found at these regions (grey: TLO), according to assembly 22. *Bottom*: Diagram depicting statistically significant enriched (red) or depleted (blue) domains for each histone modification.

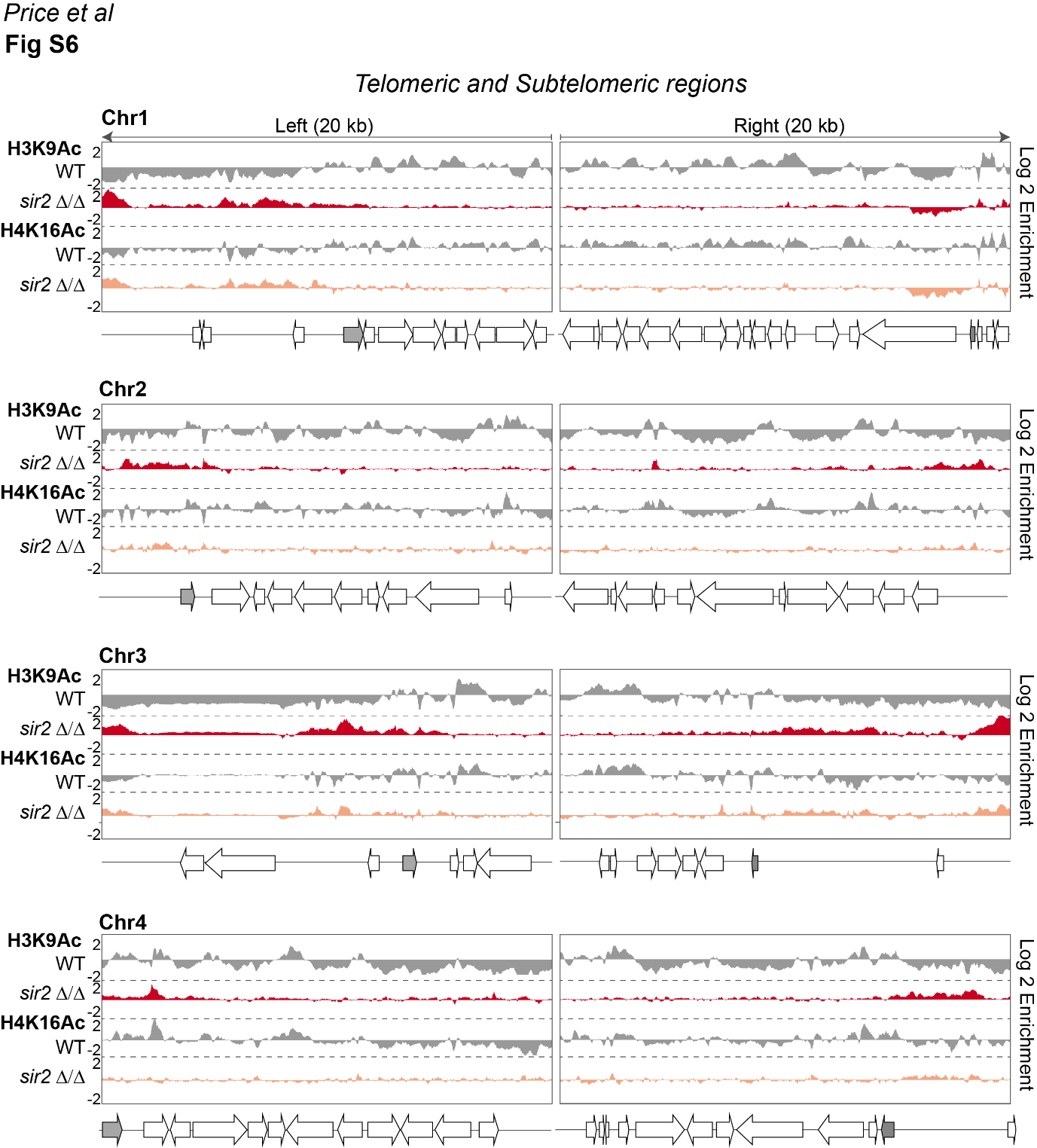

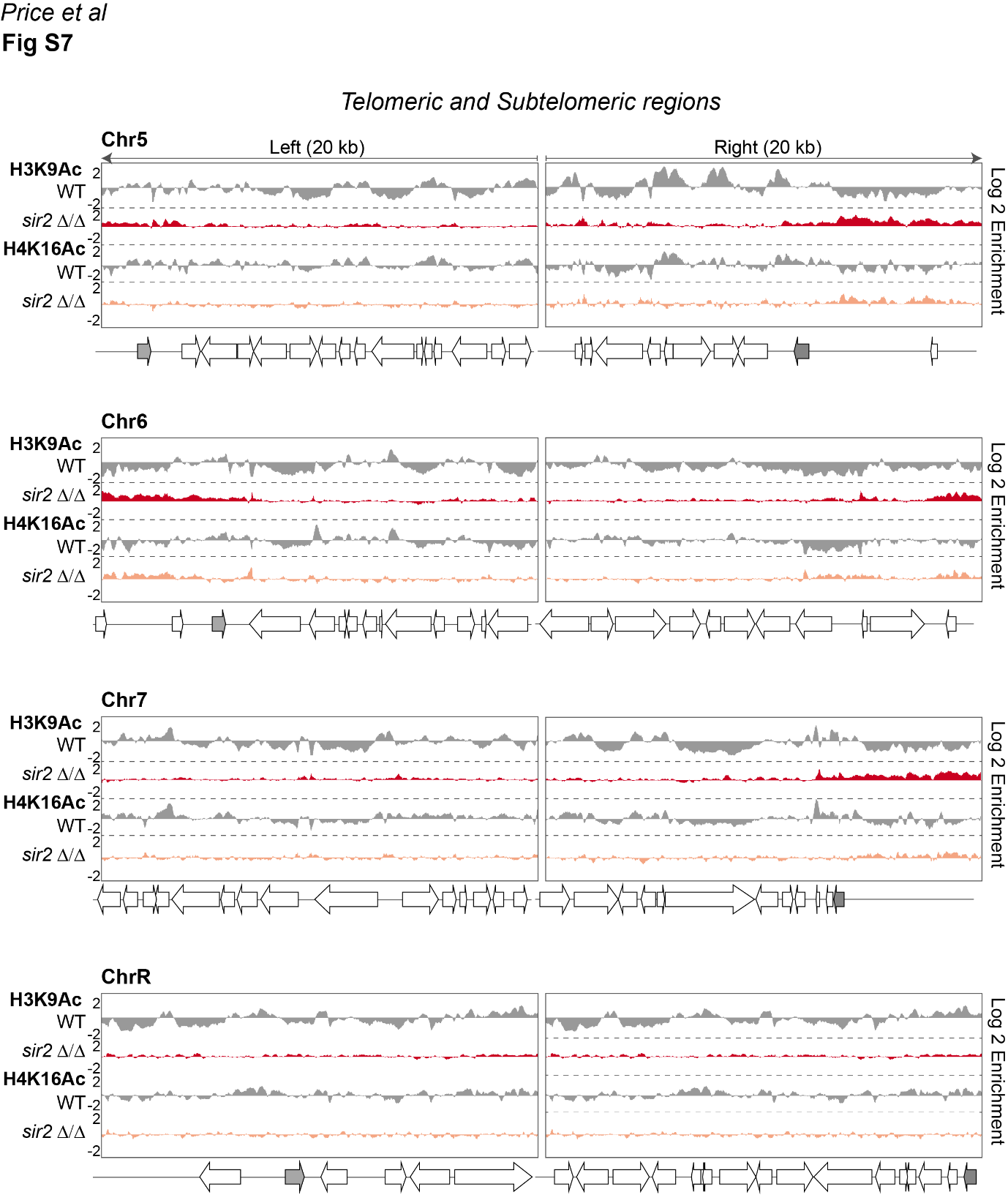

**Fig S6** and **Fig S7** *Top*: Fold enrichment (log2) of H3K9Ac and H4K16Ac relative to unmodified H4 in WT cells, and relative to WT in *sir2 Δ/Δ* cells, across the 20 kb left and right terminal regions of each of the 8 *C. albicans* chromosomes. *Bottom*: Diagrams of coding genes found at these regions (grey: TLO), according to assembly 22.

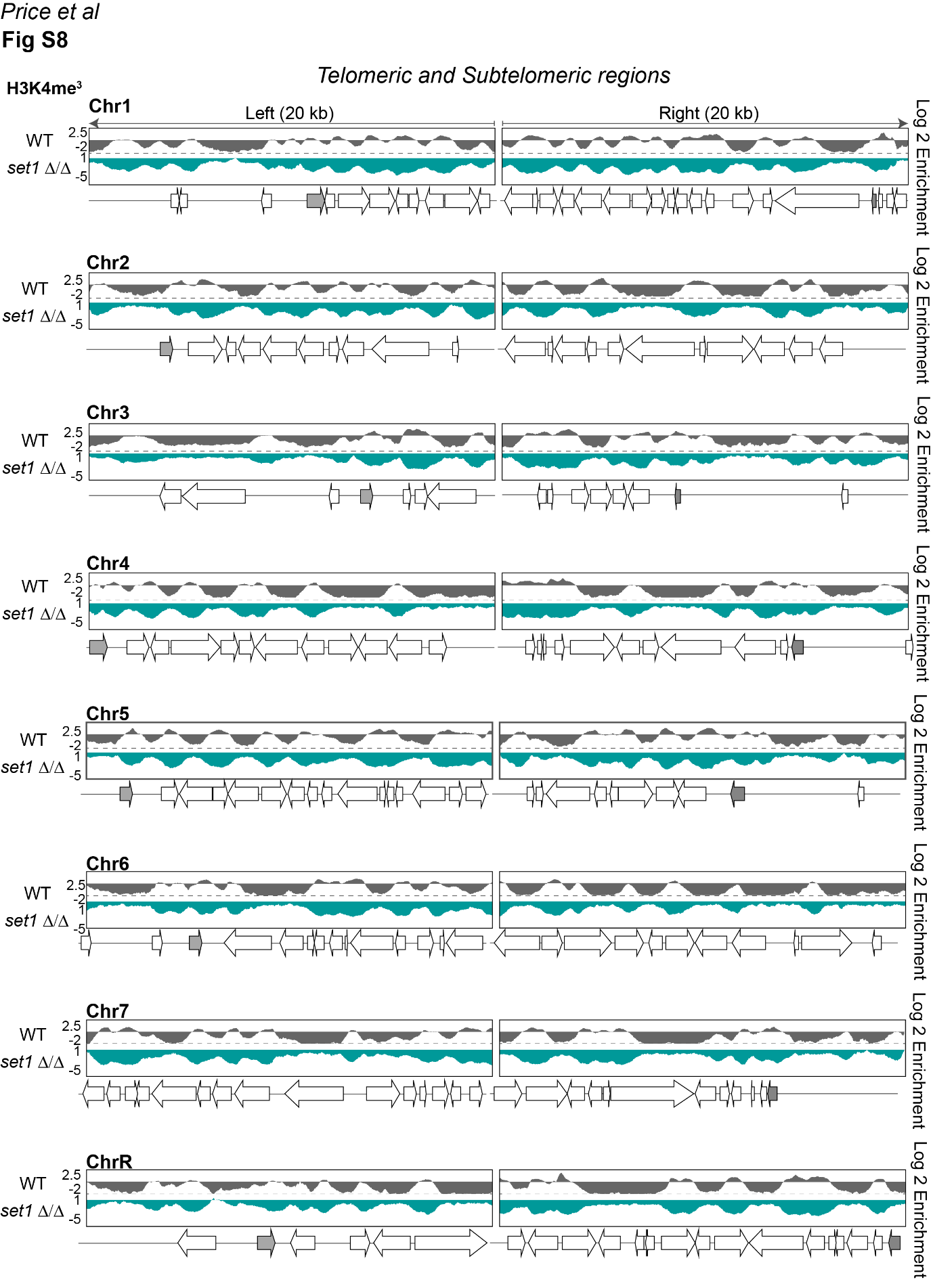

**Fig S8** *Top*: Fold enrichment (log2) of H3K4me^3^ relative to unmodified H4 in WT cells, and relative to WT in *set1 Δ/Δ* cells, across the 20 kb left and right terminal regions of each of the 8 *C. albicans* chromosomes. *Bottom*: Diagrams of coding genes found at these regions (grey: TLO), according to assembly 22.

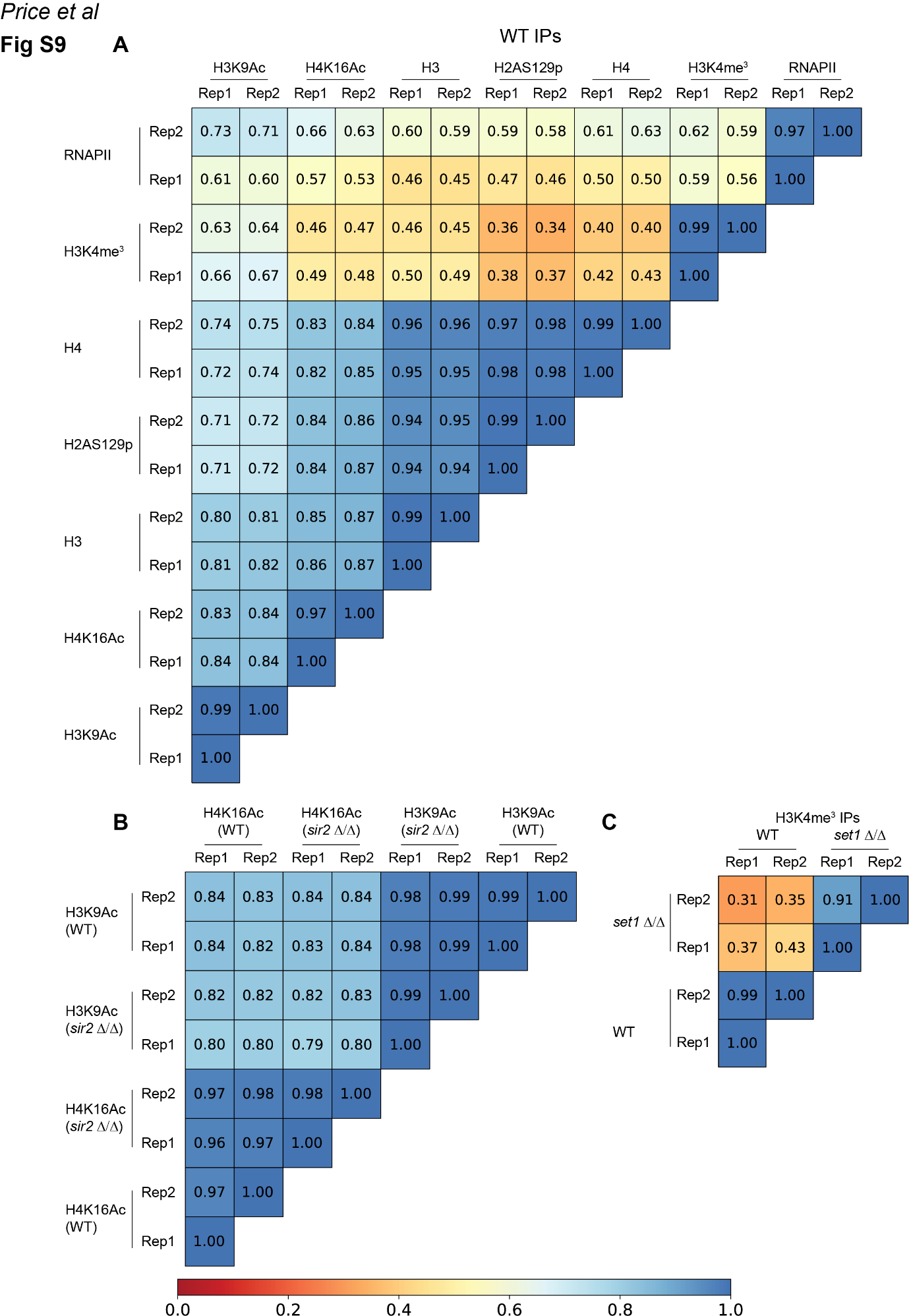

**Fig S9** Heatmaps of pairwise correlations between different IP samples. The Pearson correlation coefficients are shown. The gradient red-to-blue colour indicates the strength of correlation between the corresponding samples. **(A)** H3K9Ac, H4K16Ac, H2AS129P, H3K4me^3^, H3 and H4 IPs in WT strains; **(B)** H3K9Ac, H4K16Ac IP in WT and *sir2 Δ/Δ* strains; **(C)** H3K4me^3^ IP in WT and *set1 Δ/Δ* strains.

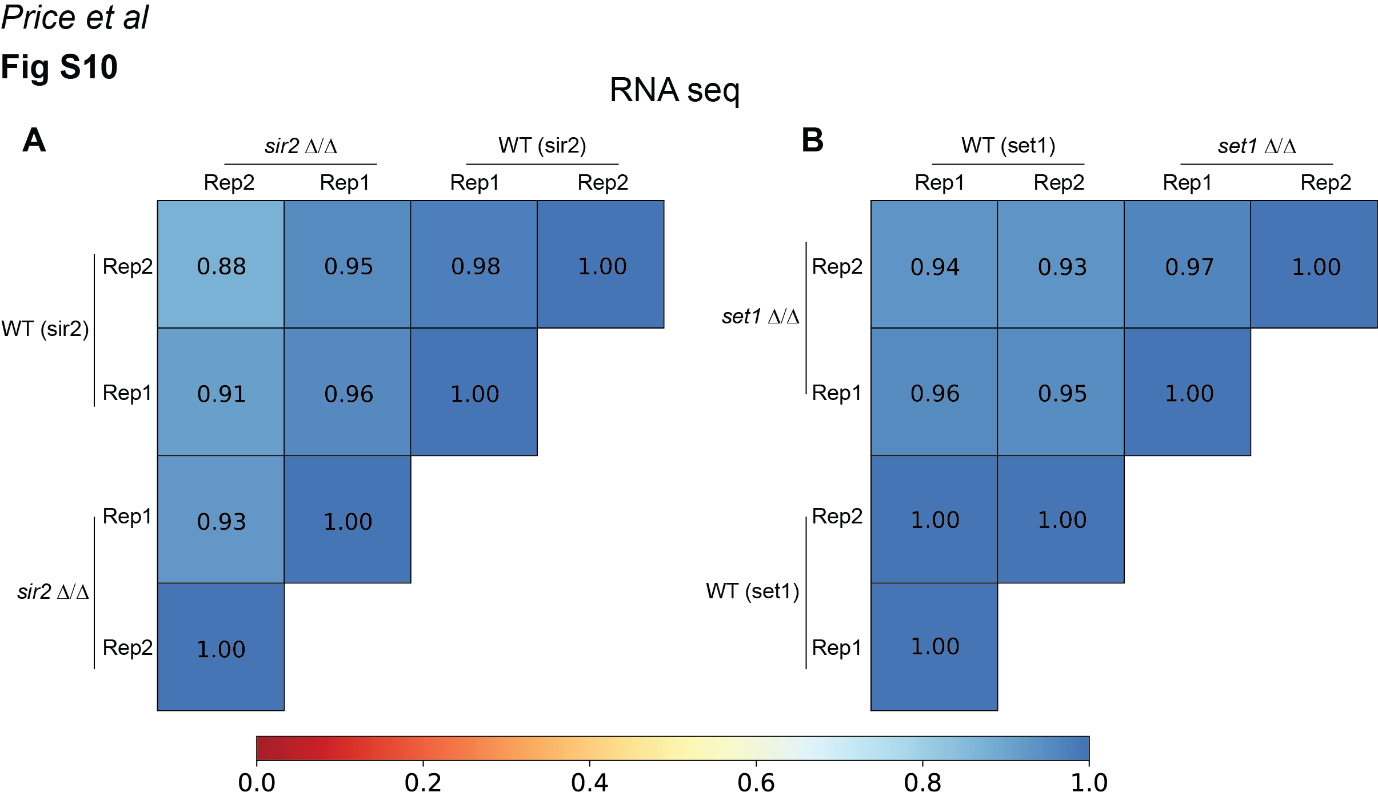

**Fig S10** Heatmap of pairwise correlations between different RNA-seq samples. The Pearson correlation coefficients are shown. The gradient red-to-blue colour indicates the strength of correlation between the corresponding samples. **(A)** RNA-seq WT and *sir2 Δ/Δ* strains; **(B)** RNA-seq in WT and *set1 Δ/Δ* strains
