## Supplementary material for "Chromatin profiling of the repetitive and non-repetitive genome of the human fungal pathogen Candida albicans"

| **Name** | **Species** | **Strain** | **Genotype** |
| --- | --- | --- | --- |
| WT (BWP17) | *C. albicans* | AB215 | ura3Δ::Δimm434/ura3Δ::Δimm434 his1::hisG/his1::hisG arg4::hisG/arg4::hisG |
| *sir2 Δ/Δ* | *C. albicans* | AB20 | ura3Δ::λimm434/ura3Δimm434 his1::hisG/his1::hisG arg4::hisG/arg4::hisG sir2::HIS1/sir2::ARG4 |
| *set1 Δ/Δ* | *C. albicans* | AB169 | ura3Δ::λimm434/ura3Δ::λimm434  his1::hisG/his1::hisG  arg4::hisG/arg4::hisG set1::HIS1/set1::ARG4 |
| WT | *S. cerevisiae* | BY4741 | MATa his3Δ1 leu2Δ0 met15Δ0 ura3Δ0 |

**Table S1**: Strains used in this study
